# Age-related changes in network controllability are mitigated by redundancy in large-scale brain networks

**DOI:** 10.1101/2023.02.17.528999

**Authors:** William Stanford, Peter J. Mucha, Eran Dayan

## Abstract

The aging brain undergoes major changes in its topology. The mechanisms by which the brain mitigates age-associated changes in topology to maintain robust control of brain networks are unknown. Here we used diffusion MRI data from cognitively intact participants (n=480, ages 40-90) to study age-associated changes in the controllability of structural brain networks, features that could mitigate these changes, and the overall effect on cognitive function. We found age-associated declines in controllability in control hubs and large-scale networks, particularly within the and frontoparietal control and default mode networks. Redundancy, quantified via the assessment of multi-step paths within networks, mitigated the effects of changes in topology on network controllability. Lastly, network controllability, redundancy, and grey matter volume each played important complementary roles in cognitive function. In sum, our results highlight the importance of redundancy for robust control of brain networks and in cognitive function in healthy-aging.

## Introduction

As populations world-wide are aging [1], dementia and other degenerative central nervous system diseases associated with cognitive decline are projected to increase in prevalence[2]. Cognitive decline is not restricted to pathological aging, but also occurs in healthy older adults. Yet healthy cognitive aging can vary greatly between individuals [3]. For those that resist cognitive decline, greater life-satisfaction, well-being, and higher levels of happiness are reported [4]. Several lifestyle factors have been found to contribute to successful cognitive aging, such as exercise [5], and education [6, 7], yet the mechanisms that could support cognitive function late in life remain incompletely understood.

Studying the topological properties of macroscopic brain connectivity with tools from network science [8] is one method by which the mechanisms that could promote cognitive function in aging were examined. Studies focused on measures of network topology that change throughout healthy [9–17], and pathological aging [18–24], and attempted to relate alterations in topology to cognition. One such central measure is network controllability [25]. Controllability is a concept that originated in engineering within the domain of control theory [26–28]. In networks, controllability examines the ability of key nodes to enable dynamic state transitions between an initial and target state [25]. For example, in brain networks the default mode network, hypothesized to be a brain state reflecting general priors for cognitive function [29], has been observed to have several hubs of average controllability [30], which are important for steering the brain towards easy to reach states. This positions the default mode network to easily direct the brain towards activity relevant for behavioral tasks [31]. Network controllability has been postulated as a dual mechanism of brain and cognitive reserve in aging by combining structural connectivity and brain dynamics to jointly measure the brain’s ability to respond and adapt to changing cognitive demands [32]. Recent studies have documented the centrality of changes in network controllability in aging [33, 34]. It nevertheless remains unknown how the brain could mitigate age-associated changes in network controllability despite changes in network topology.

A mechanism by which the brain may mitigate age-associated alterations in controllability is via increased redundancy [35–38]. Redundancy is a general principle ubiquitous in engineering that protects systems from the failure of individual components [39]. Redundancy is also evident in biological systems at many scales. Examples include at the level of genes [36, 40], organs [38], and in population coding within neural networks [41]. In the context of brain networks, redundant paths could provide alternate routes for information transmission should one path fail due to the effects of aging and/or disease. Redundant links have been previously identified as potential mechanisms that support robust control of complex networks during disconnections [25, 42, 43], but this has not been investigated in the context of network control in aging brain networks. Furthermore, redundancy has been postulated as a neuroprotective mechanism [38], but only recently studied within the context of healthy and pathological aging [12, 19, 22, 24]. It was reported that functional hippocampal redundancy supports cognitive resilience in pathological aging [19, 22, 24], and that network-wide functional redundancy mediates the relationships between age and executive function [12]. However, redundancy has yet to be investigated in the context of structural brain networks and the alterations in controllability they undergo in aging. We hypothesize that redundancy could mitigate the impact of age-associated topological changes on the controllability of brain networks.

It was recently hypothesized that network controllability and brain volume, the more traditional measure of brain reserve, should both separately be partial predictors of cognitive status [32]. In the current study we additionally attempted to investigate this aforementioned hypothesis, as well as the relevance of redundancy in structural brain networks as a potential mechanism of brain reserve. We chose processing speed as the cognitive function evaluated because it is believed to be heavily dependent on communication along white-matter tracts [44, 45]. Relatedly, processing speed is known to exhibit age-associated declines [46], that correspond with changing topological properties of structural brain networks [47, 48]. Processing speed is also associated with commonly used measures of brain reserve, such as hippocampal volume [49, 50]. We hypothesized that processing speed would be related to measures of regional influence on network dynamics, such as average controllability, particularly in functional networks that have been reported as important in age-related differences in processing speed (e.g., default mode and frontoparietal control networks [15, 51]). We expected that redundancy could support rapid communication between task-relevant functional networks [44, 45, 52], and thus be positively associated with processing speed.

To test our hypotheses we used diffusion MRI (dMRI) data from 480 participants (female = 281, male = 199) between the ages of 40-90 from the HCP-aging dataset [53] (**Fig. 1A**). We examined how average controllability, defined as the ability of brain regions to influence brain-wide dynamics, changes in aging. We constructed structural networks using the functional Schaefer local-global parcellation [54] (**Fig. 1B**). After constructing structural networks, we computed average controllability for each brain region (**Fig. 1C**), and then identified age-related shifts in average controllability of control hubs and large-scale networks (**Fig. 1C**). Next, we investigated how redundancy, a measure of multi-step paths between nodes, supports average controllability in aging (**Fig. 1D**). We hypothesized that the existence of additional pathways within structural brain networks would facilitate average controllability in networks important for cognitive function (**Fig. 1E**). Finally, we investigated the extent to which grey matter volume, network controllability, and redundancy, could serve as partial proxies of age-associated variance in processing speed (**Fig. 1F**).

**Fig. 1.**
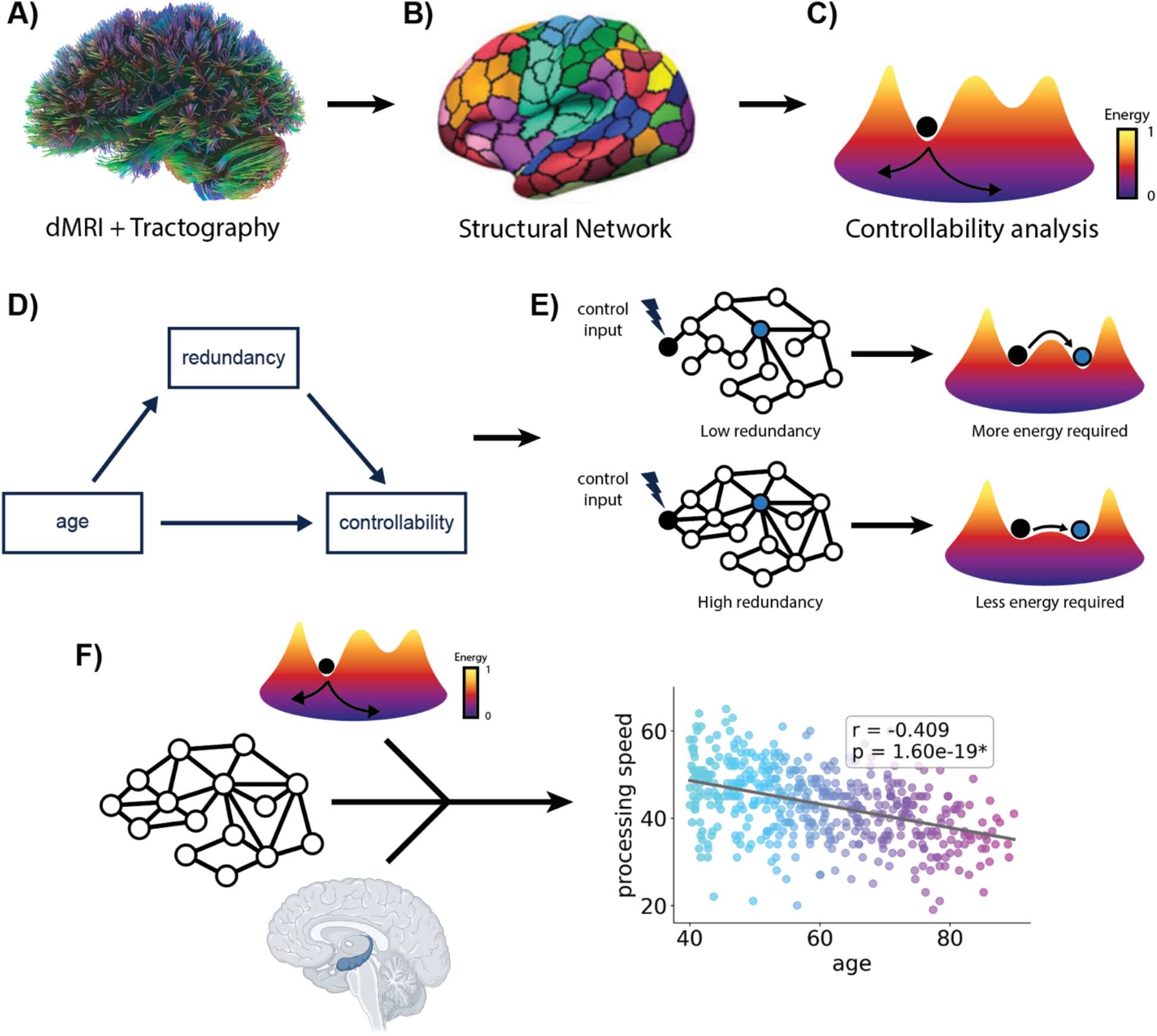
Study outline. **A)** Diffusion MRI data from 480 subjects from the HCP Aging dataset were used in our study. **B)** We constructed structural networks using the functional Schaefer local-global parcellation with 17 networks and 400 ROIs. **C)** For each subject, we calculated network controllability, a measure of a node’s ability to steer the brain into easy to reach states. **D)** We studied the relationship between controllability and network redundancy in aging, testing the extent to which redundancy influences the relationship between age and network controllability. **E)** We hypothesized that redundancy would mitigate the effects of age-associated changes in topology on average controllability in brain networks. **F)** Finally, we investigated the extent to which grey matter volume, network controllability, and redundancy, can jointly predict age-associated variance in cognitive function.

## Results

### Declines in the average controllability of control hubs and large-scale networks are implicated in aging

To begin our investigation of the relationship between network control and aging, we evaluated if the average controllability of control hubs in middle-aged participants (n = 305, ages 40-65) were different than in old-aged participants (n = 175, ages 65-90). We classified a node as a control hub if the average controllability was greater than one standard deviation above the mean average controllability for all nodes in middle-aged participants. This yielded 15 hubs, which were predominately within networks associated with cognitive function (**Fig. 2A**). The distribution of hubs within the different large-scale networks (shown as percentages in Fig. 2A) were corrected by network size by normalizing the number of hubs in each network by their respective size [30]. Hubs of average controllability were most commonly within the default mode network (~40%), followed by the salience/ventral attention network (~25%), similar to previously reported results [30]. Next, we examined if average controllability for each of these identified hubs was different between middle- and old-aged participants (**Fig. 2B**). For the hubs | with the greatest average controllability, the mean values were consistent across age_groups. However, old-aged participants had less average controllability in two hubs within the default mode network (DefaultA – PFCm_4: *F*_1, 470.75_ = 11.26, *p_bonf._* = 0.013, DefaultB – PFCd_1: *F*_1, 470.75_ = 35.01, *p_bonf._* = 9.46e-08). Next, we investigated if average controllability showed age-associated changes at the level of large-scale networks. We calculated the mean average controllability for each of the 17 large-scale networks in our parcellation, and computed the ranked Spearman’s correlation with age. Age was negatively associated with mean average controllability in the default mode (DefaultB: Spearman’s ρ = −0.303, *p_bonf._* = 2.12e-10), the frontoparietal control (ContB: Spearman’s ρ = −0.274, *p_bonf._* = 1.69e-08), and the limbic (LimbicB: Spearman’s ρ = −0.225, *p_bonf._* = 1.06e-05) networks (**Fig. 2C** and **Table S1**). For the frontoparietal control network, this decline appears to occur mostly before the age of 61 (**Fig. S1A**), whereas for the default mode and limbic networks, these declines continued throughout the age range studied (**Fig. S1B**, and **Fig S1C**, respectively).

**Fig. 2.**
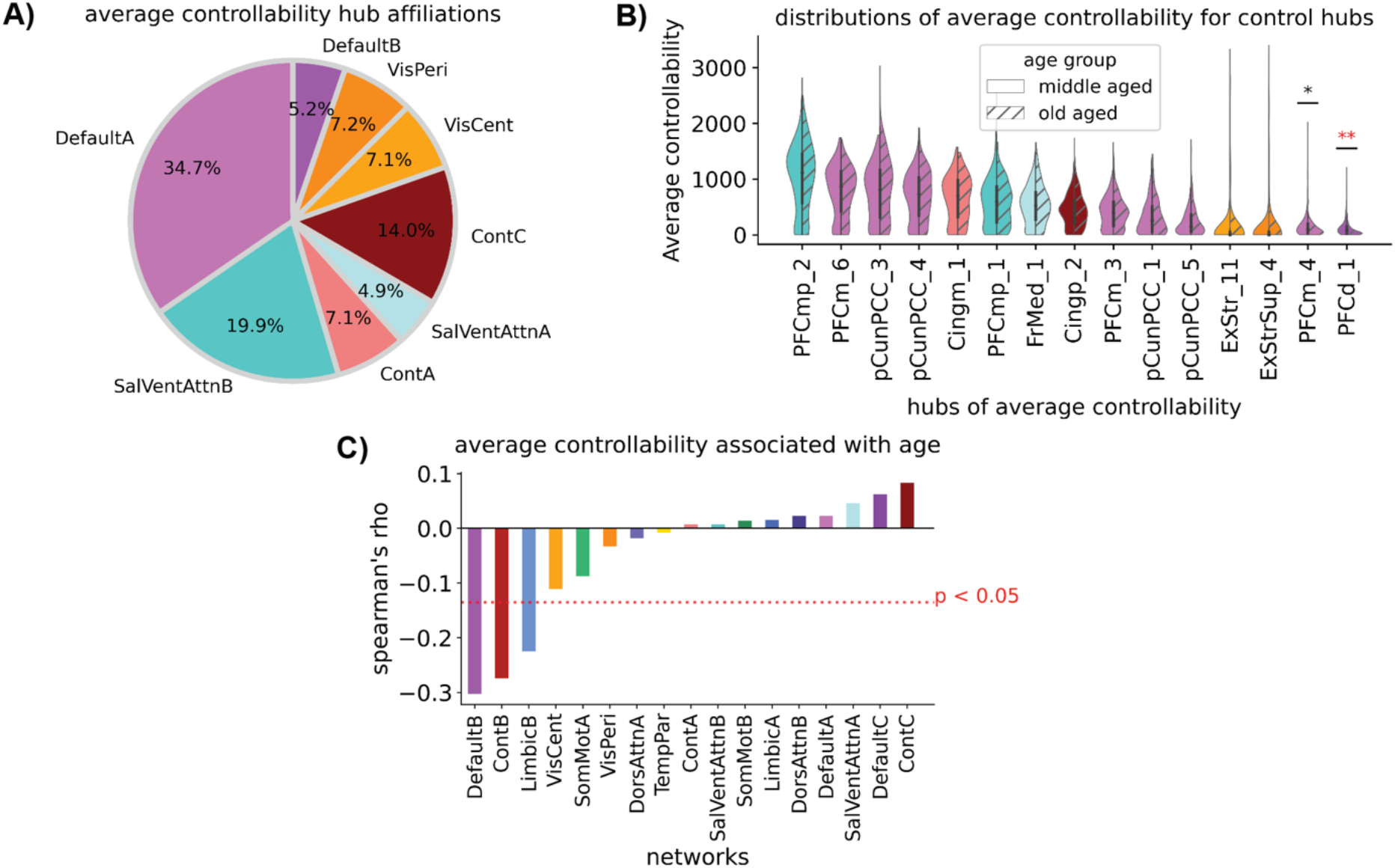
Hub and average network controllability are impacted by aging. **A)** The affiliations of hubs of average controllability in middle-aged subjects (ages 40-65) were predominately within the default mode network. Percentages were corrected by network size, which equalizes the probability of hubs falling within each network. **B)** Distributions of average controllability for each hub, for middle- and old-aged participants (ages 65-90). Two hubs in the default mode network exhibited less average controllability in old-aged participants. **C)** Average network controllability was negatively associated with age in the default mode network (DefaultA), control network (ContB), and limbic network (LimbicB). The Bonferroni method to correct for multiple comparisons was applied to correct for the number of hubs analyzed (16) (Panel B). and the number of networks (17) (Panel C). *corrected *p* < 0.05, **corrected *p* < 0.001, **corrected *p* < 1e-05.

### Degree and redundancy show similar relationships to measures of controllability

To examine if multi-step connectivity supports average controllability, we calculated network redundancy [37], defined as number of non-circular paths between nodes up to a designated length *L (see Methods*). When averaging across all subjects and performing rank correlations with average controllability, nodal degree and nodal redundancy showed similar strong positive correlations (**Fig. S2A i, iii**). Redundancy, also similarly to degree, showed a strong negative correlation with modal controllability (**Fig. S2A ii, vi**), a measure of a node’s ability to push the system to hard to reach states [55]. However, a positive relationship still existed for the rank correlation between redundancy and average controllability, adjust for degree (Spearman’s ρ = 0.444, *p* = 1.1e-20) (**Fig. S2B i**). Interestingly, the negative relationship between modal controllability and redundancy flipped to a positive correlation when regressing out the effects of ranked-degree (Spearman’s ρ = 0.388, *p* = 9.1e-16) (**Fig. 2B ii**).

### Degree mediates changes in controllability with age

Before assessing the influence of redundancy in age-associated changes in average controllability, we began by examining the importance of edges immediately connecting to nodes via degree. The average weighted degree in 13 of 17 networks showed negative rank-correlations with age (*p_bonf_* < 0.05, see **Table S2**) (**Fig. 3A**). The strongest of these associations was in the salience/ventral attention network (SalVentAttnA: Spearman’s ρ = −0.416, *p_bonf_* = 2.54e-20). With the strong relationships between degree and average controllability, we expected that average network degree should influence the association between age and average controllability. We assessed this putative relationship by testing if degree mediated age-related changes in mean network average controllability. We performed mediation analyses for each of the 17 networks, and found that degree influenced the relationship between age and average controllability for 14 of 17 networks (all *p_bonf_*’s < 0.05, see **Table S3**) (**Fig. 3B**). Degree exhibited the strongest mediation in the limbic network (LimbicB: β = −0.016,*p* < 1e-20) (**Fig. 3B**), which did not show a significant direct effect, despite significance for each other component of the mediation (**Fig. S3**).

**Fig. 3.**
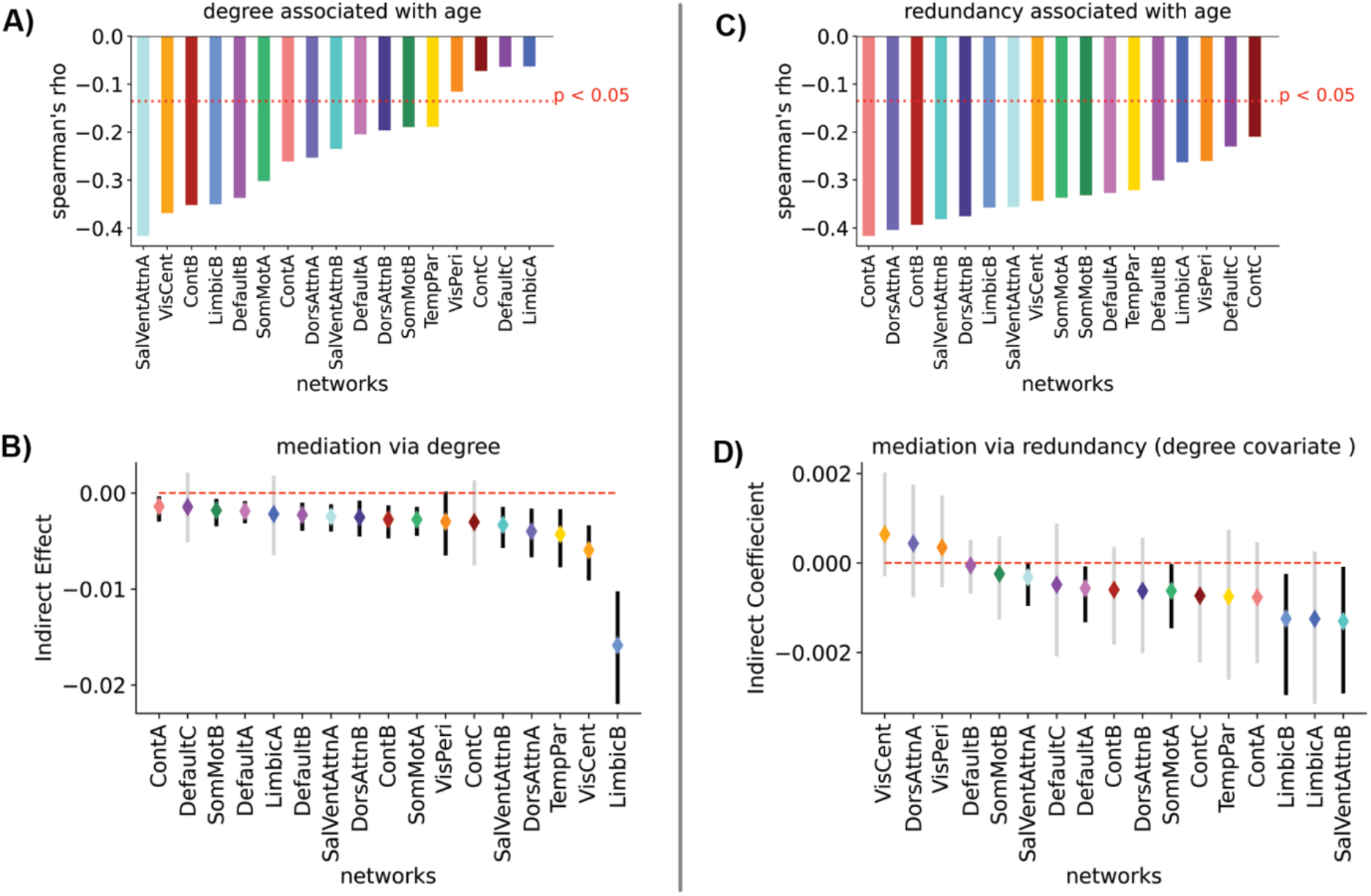
Multi-step connectivity (redundancy) mediates relationships between age and average controllability over and above the effects of degree. **A)** Average network degree was negatively associated with age for 13 of 17 large-scale networks. **B)** Changes in degree influenced the relationship between age and mean network average controllability for 14 of 17 networks. **C)** Average network redundancy also showed age associated declines, but for all networks examined. **D)** Average network redundancy mediated relationships between age and average controllability for 5 of 17 networks when including degree as a covariate. These networks included the salience/ventral attention (SalVentAttnA, SalVentAttmB), limbic (LimbicB), default mode (DefaultA), and the somatomotor (SomMotA) networks. We used the Bonferonni method to correct multiple comparisons. In each panel we corrected for the number of networks analyzed (17). In panels (B) and (D), the mediation was significant if the confidence intervals did not cross 0 when the α = 0.05/17 to correct for multiple comparisons. Significant mediations are indicated by black confidence intervals, while insignificant mediations are indicated by grey confidence intervals.

### Redundancy mediates changes in controllability with age over and above the effects of degree

We next turned towards examining the effects of redundancy in the relationship between age and average controllability. First, we calculated the average nodal redundancy for each network within our parcellation, and calculated a Spearman’s rank correlation with age. Similar to degree, average network redundancy shows widespread negative relationships with age (**Fig. 3C**). Redundancy in the frontoparietal control network should strongest relationship with age (ContA: Spearman’s *ρ* = −0.417, *p_bonf_*. = 2.24e-20), but all negative associations were significant (all *p_bonf_*’s < 0.05; see **Table S4).** Next, we investigated if the multi-step connectivity indexed by redundancy influenced the relationship between controllability and age, over and above the effects of degree. For each large-scale network in our parcellation, we performed a mediation analysis between age and average controllability with redundancy as the mediator, and included average degree of the respective large-scale networks as covariates. Redundancy mediated the relationship between age and controllability in 5 of 17 networks, over and above the effects of degree (all *p_bonf_*’s < 0.05, see **Table S5**) (**Fig. 3D**). This included the default mode network (DefaultA: *β* = −0.0006, *p_bonf_*. < 0.0004), where most average control hubs were located (**Fig. 2A**), the salient ventral attention network (SalVentAttnA: *β* = −0.0003,*p_bonf_*. < 0.0018, SalVentAttnB: *β* = −0.0013,*p_bonf_*. < 0.0010), the limbic network (LimbicB: *β* = −0.0012,*p_bonf_*. < 0.0166), which exhibited an age-associated decline in average controllability (**Fig. 2C**), and the somatomotor network (SomMotA: *β* = −0.0006, *p_bonf_*. < 0.0014). Notably, after controlling for degree, redundancy was still positively associated with average controllability in the limbic (LimbicB: *β* = 0.134,*p_bonf_*. < 0.029), salience/ventral attention (SalVentAttnB: *β* = 0.132,*p_bonf_*. < 0.032), and default mode (Default: *β* = 0.108, *p_bonf._* < 0.024) networks (**Fig. S4B**).

### Average controllability and redundancy are associated with processing speed

After focusing on age-associated variance in average controllability and network properties that contribute to it, we studied the relationships between average controllability, redundancy, and cognitive function. We first associated the mean average controllability for each large-scale network with processing speed assessed by the Pattern Comparison Processing Speed Test [56].

We hypothesized that processing speed would be related to measures of influence on overall network dynamics, such as average controllability. We found that processing speed was positively associated with average controllability in the frontoparietal control (ContB: Spearman’s *ρ* = 0.149, *p_bonf_*. = 0.019) and the default mode (DefaultB: Spearman’s *ρ* = 0.178, *p_bonf_*. = 0.002) networks (**Fig. 4A**). Next, we associated average network redundancy with processing speed. Redundancy in 4 of 17 large-scale networks was positively related to processing speed (all *p_bonf_*’s < 0.05) (**Fig. 4B**). This also included the frontoparietal control network (ContB: Spearman’s *ρ* = 0.158, *p_bonf_*. = 0.009), but redundancy in other networks was also positively associated with processing speed, including the salience/ventral attention (SalVentAttB: Spearman’s *ρ* = 0.161, *p_bonf_*. = 0.007), limbic (LimbicB: Spearman’s *ρ* = 0.158, *p_bonf_*. = 0.009), visual (VisCent: Spearman’s *ρ* = 0.148, *p_bonf_*. = 0.019), and dorsal attention (DorsAttnA: Spearman’s *ρ* = 0.148, *p_bonf_*. = 0.02) networks.

**Fig 4.**
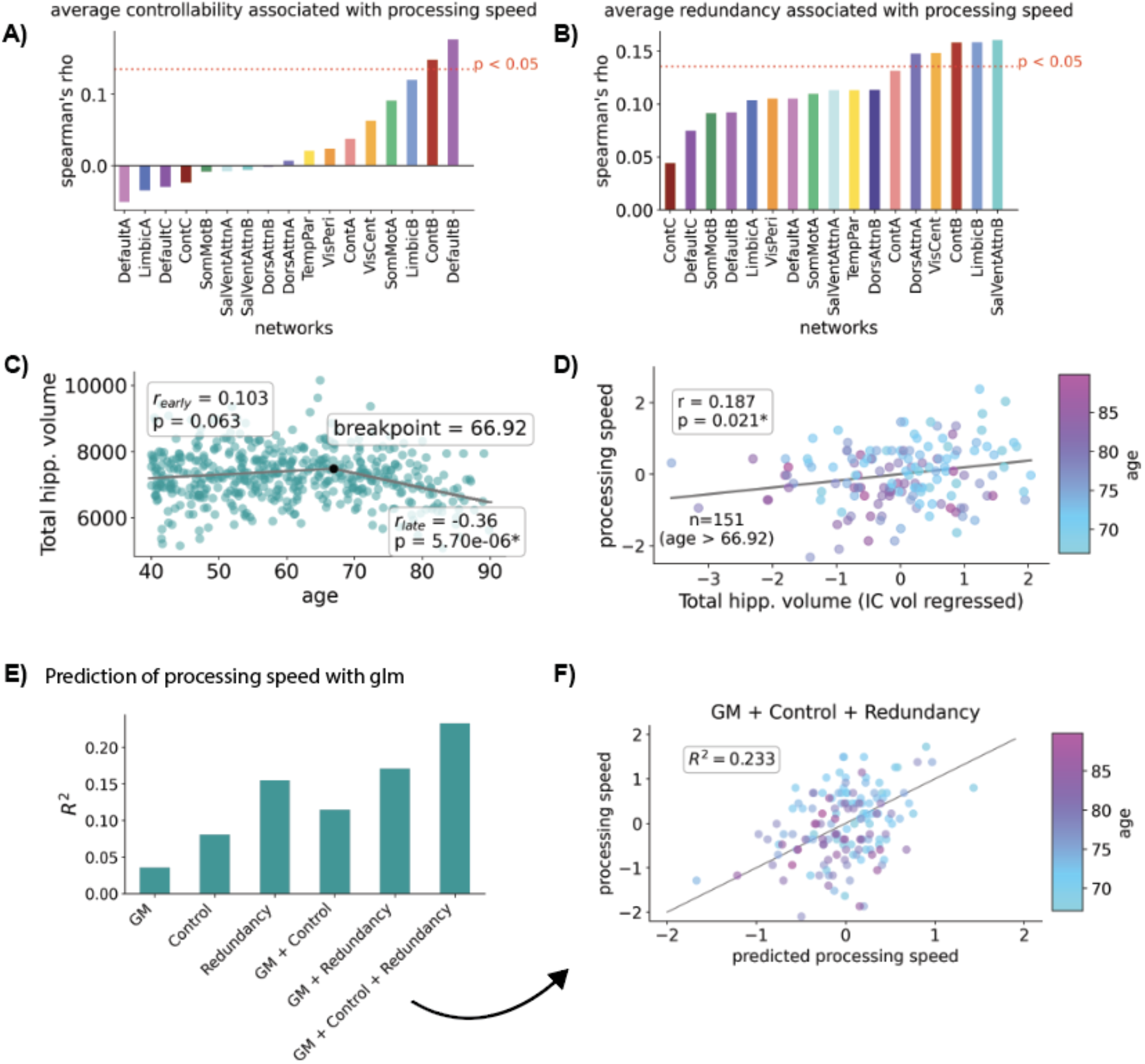
Associations between grey matter volume (GM), average controllability, and redundancy, and processing speed. **A)** Mean average controllability in the frontoparietal control (ContB), and default mode (DefaultB) networks was positively associated with processing speed. **B)** Processing speed was positively associated with redundancy in 5 of 17 networks (all *p_bonf_*’s < 0.05). **C)** Total hippocampal volume does not significantly change until around the age of 67, after which is shows a negative association with age. **D)** For participants older than 66.92, IC volume-adjusted total hippocampal volume was positively associated with processing speed. **E)** Performance of a general linear models when predicting processing speed with measures of GM volume, average controllability, and redundancy. For GM, we used IC-volume-adjusted measures of hippocampal volume, subcortical volume, and cortical volume. **F)** The z-scored predicted processing speed versus real z-scored processing speed for the best model shown in (E). In panels A, and B, we used the Bonferroni method to correct for multiple comparisons based on the number of networks analyzed (17). For panels D-E, measures of processing speed and GM volume were z-scored.

### Hippocampal grey matter volume is positively associated with processing speed in older participants

Next, we investigated the association between hippocampal grey matter (GM) volume, one of the most commonly used measures of brain reserve, and processing speed. We only expected hippocampal volume to be a mechanism of brain reserve when declines in volume began in normal aging. To determine when decline starts within our participants, we used a piece-wise linear regression that identifies breakpoints in a data-driven manner. We found that hippocampal volume experiences a non-significant but positive trend of reduction between the ages of 40 - 66.92 (*r* = 0.103, *p* = 0.063), after which hippocampal volume in our participants showed significant age-associated decline (*r* = −0.36, *p* = 5.70e-06) (**Fig. 4C**). Total subcortical GM volume showed a similar trajectory, with a breakpoint at age 65 (**Fig. S5A**), and total cortical GM volume showed continual decline throughout ages 40-90, with the most rapid decline occurring after age 75.33 (**Fig S5B**). Then, we used the residual method [57] to determine if total hippocampal volume was a marker of cognitive reserve for participants with age > 66.92. We found that total hippocampal volume, when adjusting for total intracranial volume [58], was positively associated with processing speed in this older subset of subjects (*r* = 0.187, *p* = 0.021) (**Fig. 4D**).

### Controllability, redundancy, and grey matter volume are synergistically associated with cognitive performance

Finally, we examined if network controllability, and GM volume served as complementary predictors of cognitive function in our participants. With the subset of participants older than the previously identified breakpoint (ages > 66.92), we trained GLMs to predict processing speed using various combinations of GM volume, mean average controllability, and average network redundancy for each of the 17 functional networks (**Fig. 4E**). For GM volume, we used total hippocampal volume, as well as subcortical and cortical volume. GM volume and controllability did appear to have an almost entirely complementary effect on predicting cognition, yielding an *R*^2^ = 0.115, versus an *R*^2^ = 0.036 for GM alone, and *R* = 0.081 for controllability alone. However, GM volume and redundancy showed better performance (*R*^2^ = 0.171), although there was more overlap in the predictive power between these features (redundancy: *R*^2^ = 0.155). The best model included all three sets of features (*R*^2^ = 0.233) (**Fig. 4F** and **Table 1**).

**Table 1.** GM, network controllability (Control), and network redundancy (Redundancy), each aid in the prediction of processing speed in older adults. The R^2^, log-likelihood, AIC, and BIC for each GLM trained to predict processing speed in older participants (ages > 66.92) shown in Fig 4E. Each set of features provided highly additive effects in the overall goodness-of-fit (R^2^) for these models.

| features | r<sup>2</sup> | log likelihood | AIC | BIC |
| --- | --- | --- | --- | --- |
| GM | 0.036 | -208.72 | 425.447 | -581.872 |
| Control | 0.081 | -205.1 | 446.191 | -518.647 |
| Redundancy | 0.155 | -198.87 | 433.747 | -529.614 |
| GM + Control | 0.115 | -202.33 | 446.654 | -508.628 |
| GM + Redundancy | 0.171 | -197.46 | 436.92 | -516.969 |
| Control + Redundancy | 0.218 | -193.14 | 456.279 | -453.874 |
| **GM + Control + Redundancy** | **0.233** | **-191.66** | **459.32** | **-441.154** |

## Discussion

In this study we examined whether age-associated changes in the controllability of brain networks is mitigated by redundancy. We found age-associated changes in the controllability of structural networks within our functional parcellation in the default mode (DefaultB), frontoparietal control (ContB), and limbic (LimbicB) networks. Additionally, two control hubs within the default mode network showed declines in average controllability among old-aged participants. Furthermore, we investigated the extent to which these changes were influenced by the presence of single-step and multi-step pathways between brain regions. Degree, our measure of single-step connectivity, influenced age-associated changes in average controllability in 14 of the 17 functional networks. However, multi-step paths indicative of redundancy in the system [19, 22, 24], mediated the relationships between age and average controllability in 5 of 17 networks, these included the salience/ventral attention (SalVentAttnA, SalVentAttnB), default mode (DefaultA), limbic (LimbicB), and somatomotor (SomMotA) networks. Finally, we investigated a previously posed hypothesis, that network controllability and GM volume, a more traditional measure of brain reserve, should each be partial proxies of cognitive function [32]. When using simple linear models, our results were consistent with this hypothesis. However, both redundancy and controllability appeared to provide additional predictive power when predicting the processing speed abilities of healthy older adults.

### Age related change in average controllability

Structural networks reorganize in brain aging [16]. Despite these changes, mechanisms that mitigate age-associated changes in network controllability have been relatively understudied. For modal controllability, a quantification of a brain region’s ability to push the brain into difficult to reach states [55, 59], recent work has found longitudinal changes in a multiple demand system in aging that could underly age-associated declines in executive function [33]. Other work demonstrated that the ability of temporal-parietal regions to control other brain regions decreases with age, and is particularly vulnerable to simulated lesions [34]. Our work adds to these results by evaluating changes in average controllability associated with aging in control hubs and in the structural connectivity of large-scale brain networks. We found that average controllability was similar in the top 13 of 15 control hubs between middle-aged and old-aged participants. Many of these hubs were in the precuneus, and posterior cingulate, overlapping with previously identified average control hubs [30], and regions identified as the structural core [60]. However, for two hubs in the default mode network, old-aged participants showed less average controllability than middle-aged participants. Both of these hubs were in the prefrontal cortex (PFC), one in the medial prefrontal cortex (PFCm), and the other in the dorsal prefrontal cortex (PFCd). The PFC experiences age-related declines in brain volume [61, 62] and white matter integrity [61]. Increased task-based brain activity in the PFC is commonly reported as a potential mechanism to compensate for declines in brain volume and white matter [39] (for a review see: [63, 64]). Our results suggest that increased compensatory PFC activation could also be related to declines in the average controllability of hubs within the default mode network, providing further support to the possibility of network controllability as a measure linking brain and cognitive reserve [32].

### Multi-step connectivity (redundancy) influences age-associated changes in average controllability

Nodal degree, a measure of the number edges connected to a particular node, has been shown to strongly predict nodal controllability within subjects [25, 30, 65–67]. However, the additional properties that influence controllability are largely unknown. In this study we investigated the relevance of redundant multi-step paths to brain network controllability [35–37]. We found that redundancy was positively associated with both average and modal controllability, when adjusted for degree, suggesting that multi-step pathways could play a crucial role in the control profiles of complex networks. Furthermore, our mediation analyses indicated that redundancy, while holding degree as a covariate, supported the average controllability of several key networks for cognitive function in aging. These included the salience/ventral attention, default mode (DefaultA), and limbic (LimbicB) networks. Both the default and limbic networks identified showed age-associated declines in average controllability, which indicates that redundancy could be a neuroprotective mechanism to mitigate these declines [38]. This work aligns with findings in other complex systems suggesting that edge redundancy can promote robust network controllability [25], particularly in the context of changing network topologies, such as edge removal [42, 43], which is similar to weakening white matter connectivity observed in aging [68, 69]. Our work suggests that the existence of multi-step pathways in brain networks could provide bridges of connectivity that preserve network controllability [70], to support dynamic brain activity in aging.

### Average controllability, redundancy, and processing speed

Processing speed in our study was assessed via the speed of pattern comparison [56]. This task requires several cognitive processes, such as visual search, working memory, and decision making. For both average controllability and redundancy, we found a positive association within the same subnetwork of the frontoparietal control network (ContB). The frontoparietal control network is important for allocation of attention, flexible goal-driven behavior, working memory, and decision making [71, 72]. Furthermore, the global connectivity of the frontoparietal control network may allow it to influence brain-wide dynamics [73]. Our results suggest that the average controllability, and redundancy, of edges within the frontoparietal network could be important for enabling the diverse cognitive functions relevant in processing speed and other similar tasks. Furthermore, we found that processing speed was positively associated with redundancy in the dorsal attention (DorsAttnA), visual (VisCent), and salience/ventral attention (SalVentAttnB) networks, suggesting that increased number of communication pathways involving each of these networks could support guidance of top-down attention and discrimination in this visually-based processing speed task [74–77].

### Complementary effects of grey matter volume, network controllability, and redundancy on cognitive performance

In support of the hypothesis that GM volume and network controllability could each be partial proxies of cognitive function [32], we found that GM volume and average controllability had almost directly complementary effects on the goodness of fit for our model’s prediction of processing speed. GM volume and redundancy also improved performance together versus when considering either of them alone, but the additive effect on the goodness of fit was not as dramatic. This is not surprising, as hippocampal volume has been previously associated with redundancy in brain networks [24]. Additionally, we found that average controllability and redundancy showed highly complementary effects in predicting processing speed, despite similarities in their calculation. We propose that this is primarily due to the within-subject normalization performed when calculating average controllability, which could mask age-associated variance between subjects. However, model performance was the best when including GM volume, average controllability, and redundancy in a single model, suggesting that they each could play an important role in cognitive function in healthy-aging.

### Limitations and future directions

The goals of our study included assessing the extent to which redundancy could mitigate age-associated changes in network control, and evaluate network control in the context of traditional measures of brain reserve. We used mediation analyses within our study which relied on a crosssectional sample. Cross-sectional age-associated variance does not always hold in longitudinal settings [78], thus replication of our findings in a longitudinal setting would be ideal. When studying a traditional measure of reserve, we used the residual method [57] to assess if increased hippocampal volume was positively associated with processing speed. While we did observe hippocampal atrophy (reduction in GM volume) in older participants, this method is primarily used in the context of neurodegenerative disease [57]. Future studies may consider including participants in with later stages of dementia-associated atrophy to further evaluate network controllability in the context of reserve. Lastly, the positive association between redundancy, adjusting for degree, and average controllability, as well as modal controllability, could warrant further investigation into the importance of multi-step pathways in the controllability of complex networks.

### Conclusion

In sum, we found age-associated shifts in network controllability of control hubs and large-scale brain networks in healthy middle- and old-aged adults, particularly within the default mode, frontoparietal control, and limbic networks. These age-associated changes in network controllability were mitigated by redundancy in the same networks. This suggests that, in healthy-aging, age-associated changes in network topology can be countered by the presence of redundancy in the brain.

## Methods

### Dataset and participants

We used preprocessed dMRI data obtained from the 2.0 release of the Human Connectome Project – Aging database [53]. Ages of participants ranged from 40 – 90. We restricted participants to those with normal cognitive function as assessed by the Montreal Cognitive Assessment (MoCA) [79]. For subjects older than 65, we used a cutoff of 23/30, which has been found to limit false diagnosis of mild cognitive impairment [80]. To reduce the likelihood of inclusion of participants with forms of dementia the MoCa may be less sensitive to (e.g., vascular, semantic or frontotemporal dementia), we also excluded subjects between the ages of 65-90 with poor performance on measures of cognitive flexibility, vocabulary comprehension, and executive function [17]. Poor performance was defined as a performance level worse than two standard deviations below the mean. In total, we used data from 480 (281 females, 199 males) participants for this study. All participants gave written informed consent and all procedures had been pre-approved by local Institutional Review Boards.

### Image acquisition and processing

T1-weighted structural images were acquired in a 3 Tesla Siemens Prisma Scanner. A multi-echo magnetization prepared rapid gradient echo (MPRAGE) sequence (voxel size: 0.8×0.8×0.8mm, TE = 1.8/3.6/5.4/7.2ms, TR = 2500ms, flip angle = 8 degrees) was used. Diffusion MRI (dMRI) images were generated from multi-shell diffusion with b-values of 1500 and 3000 s/mm^2^, with 93 and 92 sampling directions, a slice thickness of 1.5mm, and an in-plane resolution of 1.5mm. We used preprocessed dMRI data for our study. For details on the preprocessing pipeline see: https://brain.labsolver.org/hcp_a.html. Briefly, the pipeline involved susceptibility artifact detection with the TOPOP, from the Tiny FSL package (http://github.com/frankyeh/TinyFSL), alignment with the AC-PC line, restricted diffusion imaging [81], and generalized q-sampling [82]. These analyses were conducted at Extreme Science and Engineering Discovery Environment (XSEDE) [83] resources using the allocation TG-CIS200026.

### Network construction

Preprocessed dMRI data was reconstructed in DSI Studio (http://dsi-studio.labsolver.org). We performed whole-brain fiber tracking with 5,000,000 streamlines. (**Fig. 1A**). Structural networks were constructed according to the Schaefer Local-Global cortical parcellation with 400 cortical regions [54], which subdivides the human cortex into 17 large-scale (functional) networks (**Fig 1B**). Each brain parcel was considered a node, with the number of streamlines between any pair of parcels used as the weighted edge. A threshold of 0.001 of the maximum edge weight per subject was used as to threshold edges in the resulting brain networks.

### Average and modal controllability calculations

Average controllability, defined as the average energy from a set of control nodes on dynamic state trajectory over all possible states, was calculated using the trace of the finite time controllability Gramian [84]. The finite time controllability Gramian is computed via:

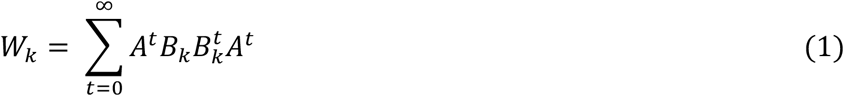

Where *A* is the adjacency matrix, and *B_k_* is an input matrix of dimension 1 *x nROIs*, and *k* represents the set of nodes specified as control nodes. For most of our analyses, we looked at the mean average controllability within each of the 17 functional networks within our parcellation, and performed rank correlations with features of interest. We also performed a supplementary analysis that involved modal controllability, defined as the ability to push the network in hard to reach states, which are the modes of the dynamical network [59]. The eigenvector of the adjacency matrix is used to compute modal controllability. Modal controllability is computed via:

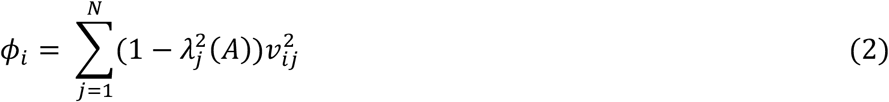

For each mode in *N*, this provides a measure of controllability from brain region *i*. In both cases, matrices were normalized dividing by one-plus the largest absolute eigenvalue before computing controllability metrics. Controllability metrics were calculated with code from: https://github.com/BassettLab/nctpy.

### Redundancy calculations

Redundancy was calculated as the number of simple (non-circular) paths between a pair of nodes up to a specified length (here we used *L* = 4) [37], according to the equation:

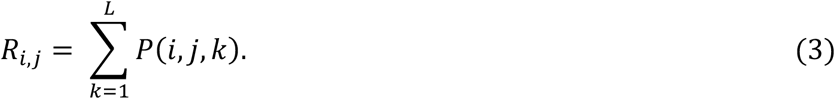

Where *P*(*i*, *j*, *k*) was the number of paths non-circular paths between nodes *i*, and *j*, calculated with the *all_simple_paths* function in NetworkX [85]. To get nodal redundancy, we summed the total number of paths from each node to all other nodes. After calculation of nodal redundancy for all nodes in each subject’s structural networks, we calculated the average redundancy in the structural connectivity of each of the 17 large-scale networks per subject. We used the binarized matrices structural connectivity matrices for these calculations.

### Cognitive measures

We focused on the cognitive measure of processing speed within our study because processing speed is believed to be limited by communication along white-matter tracts [44, 45]. Processing speed was assessed via the Pattern Comparison Processing Speed Test [56]. Subjects were shown pairs of objects and asked to judge whether two objects, presented simultaneously, were the same or different. They were given 85 seconds to judge as many objects as possible. We used participant’s MoCA scores to determine if they were healthy (score >= 23/30). Additionally, we used measures of cognitive flexibility, assessed via the used the Dimensional Card Sort Test [86], executive control, assessed via the Flanker Inhibitory Control and Attention Test [87], and vocabulary comprehension, assessed via the Picture Vocabulary Test [88], to exclude subjects with forms of dementia that the MoCA is insensitive to [17].

### Grey matter volume extraction

From the T1-weighted images, we extracted grey matter volume using the *run_first_all* command within Freesurfer [89]. This included extraction of hippocampal volume, and the volume of subcortical structures, in the Aseg atlas [90], as well cortical grey matter volume and estimated total intracranial volume.

### Statistical analysis

We performed group comparisons of average controllability in middle- and old-aged adults for control hubs. We performed rank-correlations between features of network controllability and redundancy with age and processing speed. As well as rank-correlations between redundancy, degree, and network controllability measures. We then performed mediation analyses to investigate the effects of degree and redundancy on the relationships between age and average controllability. In the mediation analysis with redundancy, ranked-degree was included as a covariate to highlight influence of multi-step pathways on the relationships between age and average controllability. Following these experiments, we performed a breakpoint analysis using a piece-wise linear regression to determine the starting point of hippocampal atrophy in our healthy cross-sectional sample. Finally, we used general linear models (GLMs) to investigate the extent to which linear combinations of GM and network features aided in the prediction of processing speed. Welch’s ANOVAs, Spearman’s correlations, Pearson’s correlations, and mediation analysis, were performed using the python package Pingouin [91]. For the Welch’s ANOVAs, we compared average controllability in 15 identified hubs in middle- and old-aged adults. We used the Bonferroni method to correct for multiple comparisons which set the p-value necessary for significance to p < 0.05/15. For Spearman and Pearson correlations, we required p < 0.05/17 to correct for the number of functional networks analyzed. Mediation analyses were performed with the *α* = 0.05/17 for the confidence intervals, with significance determined by whether or not the confidence intervals for each coefficient crossed the value of zero. Piece-wise linear regression to determine breakpoints in rates of change for grey matter volume was performed using the pwlf python package [92]. Our GLMs were constructed using the python package Statsmodels [93]. Additional stats derived from these models (*R*^2^, log-likelihood, AIC [94], BIC [95]) were also computed using the Statsmodels package.

### Plotting

We used custom python scripts for plotting and data visualization based on the Matplotlib [96], Pandas [97], and Seaborn [98], packages.

## Supporting information

Stanford-bioRxiv-23-SM

## Acknowledgments

We thank Dr. Kelly Giovanello for helpful suggestions.

## Author contributions

Conceptualization: WS, PJM, ED

Methodology: WS

Investigation: WS

Visualization: WS

Supervision: PJM, ED

Writing, review & editing: WS, PJM, ED

## Author’s note

Peter J. Mucha is currently serving as an Associate Editor on the Social and Interdisciplinary Sciences section at Science Advances.

## Data and materials availability

All data used in this study is publicly available via the Human Connectome Project - Aging dataset [53]. All reported results are available in the main text or in the supplementary materials.

