## Supplementary material for "Age-related changes in network controllability are mitigated by redundancy in large-scale brain networks": Stanford-bioRxiv-23-SM

William Stanford *et al.*

**This PDF file includes:**

Figs. S1 to S5
Tables S1 to S5

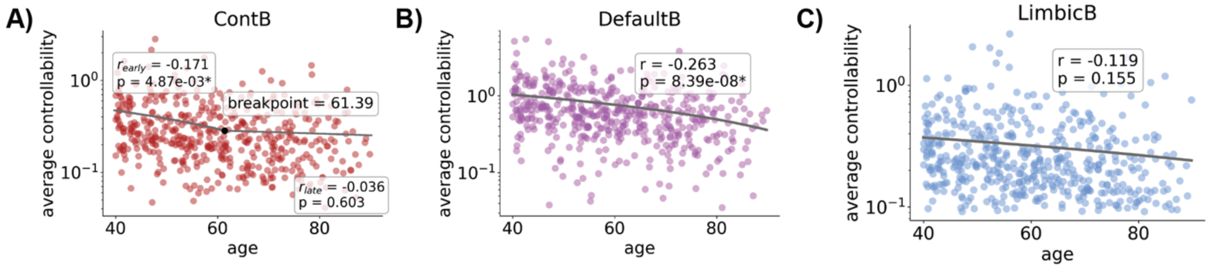


**Fig. S1. Scatter plots of age versus the mean average controllability for networks that showed negative rank correlations with age.** For the purpose of illustrating the raw values, we report Pearson correlations here, whereas the rank correlations are reported in the main paper. **A)** The frontoparietal control network (ContB) only showed decline between the ages of 40-61, afterwards the rate of decline was not significant from zero. **B)** The default mode network (DefaultB) showed a negative relationship with age throughout the entire age-range studied. **C)** The limbic network (LimbicB), did not show a significant linear correlation, despite the significant rank correlation observed in the main text.

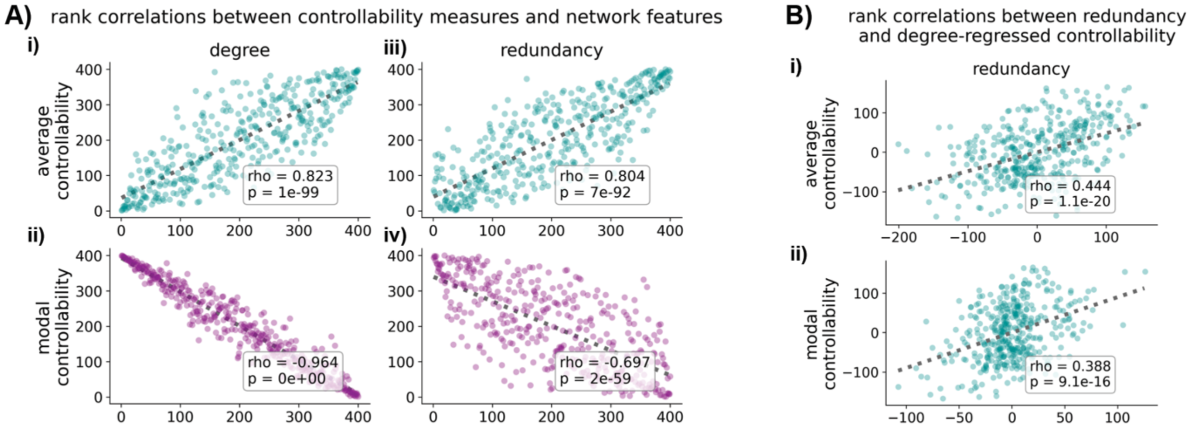


**Fig. S2. Redundancy relates to measures of controllability over and above the effects of degree. A)** Degree and redundancy showed similar relationships with measures of average controllability and modal controllability. **B)** Degree adjusted redundancy was still positively associated with average controllability (i). For modal controllability the relationship with modal controllability was flipped when regressing out the effects of degree (ii).

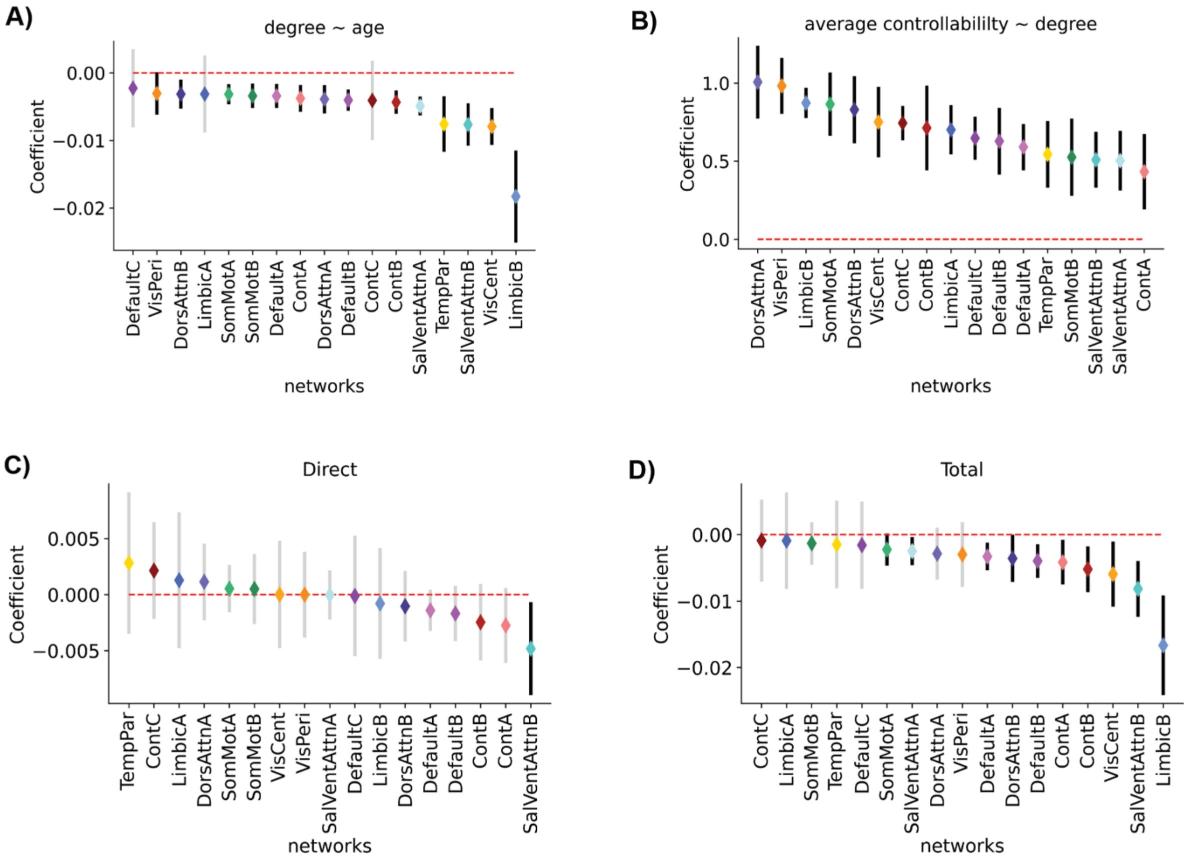


**Sig S3. The relationships between degree and age, average controllability and degree, and direct and total effects for the mediation analysis presented in Fig 3B. A)** Degree was negatively related to participant age in 14 of 17 networks. **B)** Average controllability was positively related to degree in all networks. **C)** There was a negative direct relationship between age and average controllability in the salience/ventral attention network (SalVentAttnB). **D)** The total effect was significant in 10 of 17 networks. Significance was determined if the confidence intervals for each coefficient did not cross zero after setting the $\alpha = 0.05/17$ to correct for multiple comparisons.

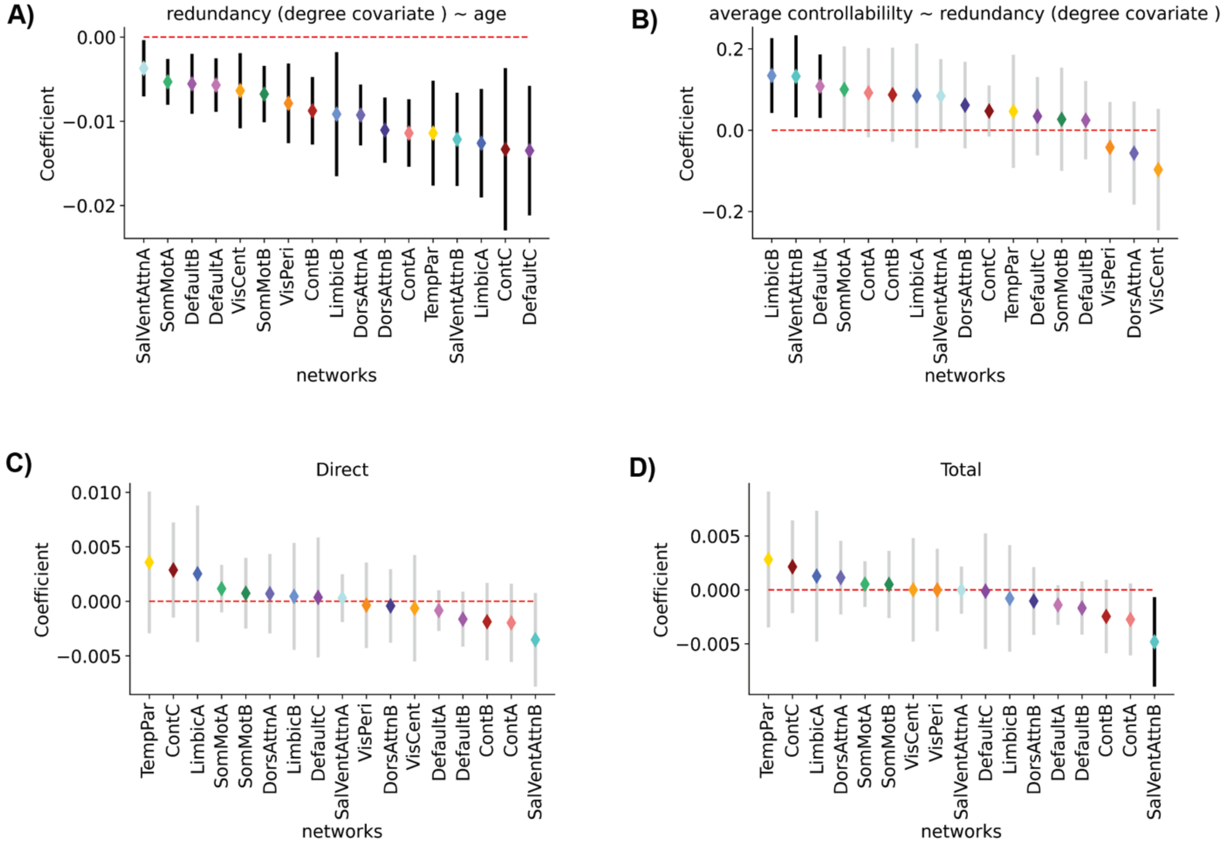


**Fig S4. The relationships between redundancy and age, average controllability and redundancy, and direct and total effects for the mediation analysis presented in Fig 3D.** Degree was included as a covariate in this analysis to examine the effects over and above the effect of degree. **A)** Redundancy was negatively related to participant age in all networks. **B)** Average controllability was positively related to degree in the limbic (LimbicB), salience/ventral attention (SalVentAttnB), and default mode (DefaultA) networks. **C)** No significant negative direct relationship between age and average controllability existed when controlling for degree. **D)** There was a single significant total effect within the salience/ventral attention network (SalVentAttnB). Significance was determined if the confidence intervals for each coefficient did not cross zero after setting the $\alpha = 0.05/17$ to correct for multiple comparisons.

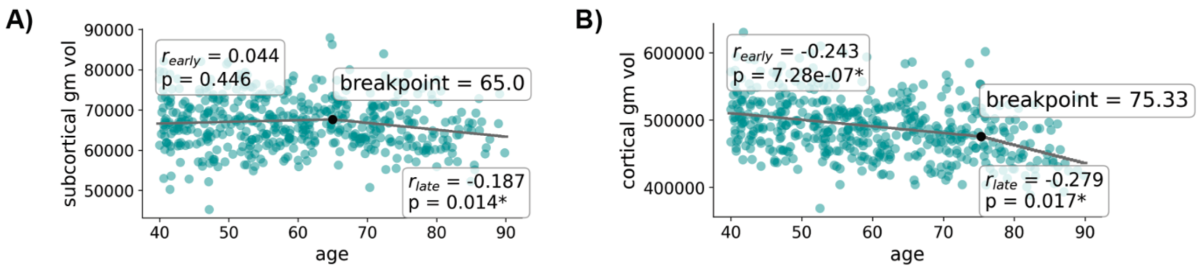


**Fig S5. Age-associated changes in subcortical and cortical grey matter volume. A)** Subcortical grey matter volume showed no age-associated declines until age 65, afterwards it was negatively associated with age. **B)** Cortical grey matter volume experienced age-associated decline across the entire age-range studied, but declined more quickly after the age of ~75. We used a piece-wise regression that determines the break-point in a data-driven manner.

**Supplementary tables S1-S6**

Average network average controllability associated with age

| Network | $p_{bonf.}$ | spearman's ρ | CI95%_l | CI95%_u |
|---|---|---|---|---|
| DefaultB* | 2.12E-10 | -0.303 | -0.38 | -0.22 |
| ContB* | 1.69E-08 | -0.274 | -0.36 | -0.19 |
| LimbicB* | 1.06E-05 | -0.225 | -0.31 | -0.14 |
| VisCent | 0.25 | -0.111 | -0.2 | -0.02 |
| SomMotA | 0.933 | -0.088 | -0.18 | 0 |
| VisPeri | 0.995 | -0.033 | -0.12 | 0.06 |
| DorsAttnA | 0.995 | -0.018 | -0.11 | 0.07 |
| TempPar | 0.995 | -0.008 | -0.1 | 0.08 |
| ContA | 0.995 | 0.007 | -0.08 | 0.1 |
| SalVentAttnB | 0.995 | 0.007 | -0.08 | 0.1 |
| SomMotB | 0.995 | 0.014 | -0.08 | 0.1 |
| LimbicA | 0.995 | 0.015 | -0.07 | 0.1 |
| DorsAttnB | 0.995 | 0.022 | -0.07 | 0.11 |
| DefaultA | 0.995 | 0.022 | -0.07 | 0.11 |
| SalVentAttnA | 0.995 | 0.046 | -0.04 | 0.13 |
| DefaultC | 0.995 | 0.062 | -0.03 | 0.15 |
| ContC | 0.995 | 0.083 | -0.01 | 0.17 |

**Table S1. The statistics and confidence intervals for rank correlations of average network average controllability with age.** Networks are sorted in ascending order by the calculated Spearman's ρ's. The Bonferoni method was used to correct for multiple comparisons. (*$p_{bonf} < 0.05$)

| Average network degree associated with age | | | | |
|---|---|---|---|---|
| **Network** | ***p* bonf.** | **spearman's ρ** | **CI95%_l** | **CI95%_u** |
| SalVentAttnA* | 2.54E-20 | -0.416 | -0.49 | -0.34 |
| VisCent* | 1.06E-15 | -0.369 | -0.44 | -0.29 |
| ContB* | 3.40E-14 | -0.352 | -0.43 | -0.27 |
| LimbicB* | 4.58E-14 | -0.35 | -0.43 | -0.27 |
| DefaultB* | 5.32E-13 | -0.337 | -0.41 | -0.26 |
| SomMotA* | 2.39E-10 | -0.302 | -0.38 | -0.22 |
| ContA* | 1.07E-07 | -0.261 | -0.34 | -0.18 |
| DorsAttnA* | 3.24E-07 | -0.253 | -0.33 | -0.17 |
| SalVentAttnB* | 3.11E-06 | -0.235 | -0.32 | -0.15 |
| DefaultA* | 1.06E-04 | -0.205 | -0.29 | -0.12 |
| DorsAttnB* | 2.40E-04 | -0.197 | -0.28 | -0.11 |
| SomMotB* | 4.84E-04 | -0.19 | -0.27 | -0.1 |
| TempPar* | 5.02E-04 | -0.189 | -0.27 | -0.1 |
| VisPeri | 0.189 | -0.116 | -0.2 | -0.03 |
| ContC | 0.995 | -0.073 | -0.16 | 0.02 |
| DefaultC | 0.995 | -0.064 | -0.15 | 0.03 |
| LimbicA | 0.995 | -0.063 | -0.15 | 0.03 |

**Table S2. The statistics and confidence intervals for rank correlations of average network degree with age.** Networks are sorted in ascending order by the calculated Spearman's ρ's. The Bonferoni method was used to correct for multiple comparisons. (*$p_{bonf} < 0.05$).

| Indirect effects of degree on the relationships between age and average controllability | | | | | | |
|---|---|---|---|---|---|---|
| **Network** | **coef** | **se** | **pval** | **CI[0.1%]** | **CI[99.9%]** | **sig** |
| ContA | -0.001 | 0.000 | 0.000 | -0.003 | -0.00055 | Yes |
| DefaultC | -0.001 | 0.001 | 0.221 | -0.005 | 0.0019538 | No |
| SomMotB | -0.002 | 0.000 | 0.000 | -0.003 | -0.000821 | Yes |
| DefaultA | -0.002 | 0.000 | 0.000 | -0.003 | -0.001046 | Yes |
| LimbicA | -0.002 | 0.001 | 0.082 | -0.006 | 0.00161 | No |
| DefaultB | -0.002 | 0.000 | 0.000 | -0.004 | -0.00122 | Yes |
| SalVentAttnA | -0.002 | 0.000 | 0.000 | -0.004 | -0.001387 | Yes |
| DorsAttnB | -0.003 | 0.001 | 0.000 | -0.004 | -0.001013 | Yes |
| ContB | -0.003 | 0.000 | 0.000 | -0.005 | -0.001504 | Yes |
| SomMotA | -0.003 | 0.000 | 0.000 | -0.004 | -0.001662 | Yes |
| VisPeri | -0.003 | 0.001 | 0.003 | -0.006 | -4.97E-05 | Yes |
| ContC | -0.003 | 0.001 | 0.033 | -0.007 | 0.0011253 | No |
| SalVentAttnB | -0.003 | 0.001 | 0.000 | -0.006 | -0.001671 | Yes |
| DorsAttnA | -0.004 | 0.001 | 0.000 | -0.006 | -0.001842 | Yes |
| TempPar | -0.004 | 0.001 | 0.000 | -0.008 | -0.001888 | Yes |
| VisCent | -0.006 | 0.001 | 0.000 | -0.009 | -0.003583 | Yes |
| LimbicB | -0.016 | 0.002 | 0.000 | -0.022 | -0.010454 | Yes |

**Table S3. Statistics for the mediations of degree on the relationship between age and average controllability for each of the 17 networks.** For all but 3 networks there was a significant mediation by degree ($p_{bonf.} < 0.05$). Significance was determined if the confidence intervals for each coefficient did not cross zero after setting the $\alpha = 0.05/17$ to correct for multiple comparisons.

Average network redundancy associated with age

| Network | $p_{bonf.}$ | spearman's ρ | CI95%_l | CI95%_u |
|---|---|---|---|---|
| ContA* | 2.24E-20 | -0.417 | -0.49 | -0.34 |
| DorsAttnA* | 4.37E-19 | -0.404 | -0.48 | -0.33 |
| ContB* | 4.84E-18 | -0.394 | -0.47 | -0.32 |
| SalVentAttnB* | 6.714E-17 | -0.382 | -0.46 | -0.3 |
| DorsAttnB* | 2.603E-16 | -0.376 | -0.45 | -0.3 |
| LimbicB* | 1.038E-14 | -0.358 | -0.43 | -0.28 |
| SalVentAttnA* | 1.282E-14 | -0.357 | -0.43 | -0.28 |
| VisCent* | 1.451E-13 | -0.344 | -0.42 | -0.26 |
| SomMotA* | 5.026E-13 | -0.338 | -0.41 | -0.26 |
| SomMotB* | 1.385E-12 | -0.332 | -0.41 | -0.25 |
| DefaultA* | 3.155E-12 | -0.327 | -0.41 | -0.25 |
| TempPar* | 8.773E-12 | -0.322 | -0.4 | -0.24 |
| DefaultB* | 2.715E-10 | -0.301 | -0.38 | -0.22 |
| LimbicA* | 7.995E-08 | -0.263 | -0.34 | -0.18 |
| VisPeri* | 1.161E-07 | -0.261 | -0.34 | -0.18 |
| DefaultC* | 5.62E-05 | -0.23 | -0.31 | -0.14 |
| ContC* | 6.60E-05 | -0.21 | -0.29 | -0.12 |

**Table S4. The statistics and confidence intervals for rank correlations of average network redundancy with age.** Networks are sorted in ascending order by the calculated Spearman's ρ's. The Bonferoni method was used to correct for multiple comparisons. (*$p_{bonf}$< 0.05)

Indirect effects of redundancy on the relationships between age and average controllability

| Network | coef | se | pval | CI[0.1%] | CI[99.9%] | sig |
|---|---|---|---|---|---|---|
| VisCent | 0.0006 | 0.0004 | 0.0328 | -0.0002 | 0.0020 | No |
| DorsAttnA | 0.0004 | 0.0004 | 0.2524 | -0.0007 | 0.0017 | No |
| VisPeri | 0.0003 | 0.0003 | 0.2130 | -0.0005 | 0.0015 | No |
| DefaultB | -0.0001 | 0.0002 | 0.7390 | -0.0006 | 0.0005 | No |
| SomMotB | -0.0002 | 0.0003 | 0.3692 | -0.0012 | 0.0006 | No |
| SalVentAttnA | -0.0003 | 0.0001 | 0.0018 | -0.0009 | 0.0000 | Yes |
| DefaultC | -0.0005 | 0.0004 | 0.2854 | -0.0020 | 0.0008 | No |
| DefaultA | -0.0006 | 0.0002 | 0.0004 | -0.0013 | -0.0001 | Yes |
| ContB | -0.0006 | 0.0003 | 0.0626 | -0.0018 | 0.0003 | No |
| DorsAttnB | -0.0006 | 0.0004 | 0.1208 | -0.0020 | 0.0005 | No |
| SomMotA | -0.0006 | 0.0002 | 0.0014 | -0.0014 | -0.0001 | Yes |
| ContC | -0.0007 | 0.0003 | 0.0076 | -0.0022 | 0.0000 | No |
| TempPar | -0.0008 | 0.0005 | 0.1370 | -0.0026 | 0.0007 | No |
| ContA | -0.0008 | 0.0004 | 0.0604 | -0.0022 | 0.0004 | No |
| LimbicB | -0.0012 | 0.0004 | 0.0002 | -0.0029 | -0.0003 | Yes |
| LimbicA | -0.0012 | 0.0006 | 0.0166 | -0.0031 | 0.0002 | No |
| SalVentAttnB | -0.0013 | 0.0004 | 0.0010 | -0.0029 | -0.0001 | Yes |

**Table S5. Statistics for the mediations of redundancy with degree as a covariate on the relationship between age and average controllability for each of the 17 networks.** For all but 5 of 17 networks there was a significant mediation by redundancy ($p_{bonf.} < 0.05$). Significance was determined if the confidence intervals for each coefficient did not cross zero after setting the $\alpha = 0.05/17$ to correct for multiple comparisons.
